# Impact of Absent in Melanoma 2 (AIM2) on Antigen-Specific CD4+ T cell Activation and Homeostasis

**DOI:** 10.64898/2025.12.30.697014

**Authors:** Dakota M. Reinartz, Chloe Sairs, Wang Gong, Yu Leo Lei, Michael S. Kuhns, Justin E. Wilson

**Affiliations:** Department of Immunobiology, University of Arizona, Tucson, AZ; University of Arizona Cancer Center, Tucson, AZ; Department of Cancer Biology, University of Texas M.D. Anderson Cancer Center, Houston, TX; Department of Head and Neck Surgery, University of Texas M.D. Anderson Cancer Center, Houston, TX; Cancer Biology Graduate Interdisciplinary Program, University of Arizona, Tucson, AZ; The BIO-5 Institute, University of Arizona, Tucson, AZ; The Arizona Center on Aging, The University of Arizona College of Medicine, Tucson, AZ

## Abstract

The intrinsic role of Absent in Melanoma 2 (AIM2) in CD4*+* T cells during antigen-specific activation and differentiation is not fully understood. To address this, we crossed AIM2-deficient mice with OT-II/RAG transgenic mice, which express an ovalbumin-specific T cell receptor in CD4*+* T cells (OT-II). We found that AIM2 does not regulate thymic selection of CD4*+* thymocytes, but may promote trafficking or survivability of CD4*+* T cells in the spleen. In vitro coculture assays revealed that *Aim2^-/-^* OT-II CD4*+* T cells produce significantly less IL-2 than wild type OT-II CD4*+* T cells when cocultured with ovalbumin-incubated dendritic cells. However, we found no differences in CD4*+* T cell proliferative capacity and differentiation among WT OT-II and *Aim2^-/-^* OT-II CD4*+* T cells following OVA immunization in vivo. Finally, adoptively transferred WT and *Aim2^-/-^* OT-II cells controlled the growth of implanted OVA-expressing NOOC2 syngeneic tumors at an equal capacity. Taken together, these results indicate that AIM2 contributes to CD4*+* T cell homeostasis in secondary lymphoid organs in vivo. AIM2 also intrinsically promotes IL-2 production in response to antigen-specific activation in vitro. However, this loss of IL-2 does not translate to measurable defects in proliferative, differentiation or effector function during OVA immunization or tumor challenge in vivo.

## Introduction

T cells play crucial roles in balancing adaptive immune responses against a variety of immunological insults and developing tolerance. CD4^+^ T cells recognize peptide antigens presented by class II MHC molecules and differentiate into one of several T helper cell subsets primarily based on the TCR signal strength [1] along with the cytokine environment it encounters during antigen-MHC engagement. These CD4 subsets, include Th1, Th2, Th17, T follicular helper (Tfh) cells and regulatory T cells (Tregs) [2], which function to direct the response of other immune cells towards appropriate immunity or tolerance against the antigenic source that activated the T cell. T cells require a variety of molecular mechanisms to become appropriately activated and differentiated. Thus, it is crucial to understand the molecular that guide these processes.

For a CD4^+^ T cell to fully activate, it must receive three major signals. This includes: 1) engagement of T cell receptor (TCR), CD3 and CD4 with peptide-MHC class II expressed on the surface of an antigen presenting cell (APC); 2) co-stimulation through engagement of T cell-bound CD28 to CD80/CD86 expressed by APCs; and 3) signaling via cytokines, which further aides in directing CD4^+^ T cell proliferation and differentiation. Among these is IL-2, a pleiotropic cytokine produced by T cells that can act in an autocrine fashion to regulate clonal expansion and the expression of master transcription factors that determine T cell fates [3]. This includes the promotion of T-bet, the lineage defining transcription factor for Th1 cells, and inhibition of BCL6, the lineage defining transcription factor for Tfh cells. In coordination with TGF-β, IL-2 commits a CD4^+^ T cell towards the Treg lineage [4]. T cell differentiation and establishment of immunological memory is further influenced by additional cytokines and environmental factors encountered during activation. Cellular damage-associated molecular patterns (DAMPs) have recently been implicated during T cell activation and differentiation [5] as well driving the establishment of memory populations within tissue [6], however the sensors utilized by T cells during DAMP recognition have not been fully characterized.

Absent in Melanoma 2 (AIM2) is a pattern recognition receptor that binds to cytosolic double-stranded DNA (dsDNA) derived from pathogens and the host. DNA binding triggers formation of the AIM2 inflammasome which leads to activation of Caspase-1 and release of mature IL-1β and IL-18 and in some cases, an inflammatory form of cell death termed pyroptosis [7]. AIM2 also performs inflammasome-independent functions. This includes suppressing AKT phosphorylation, which is vital to metabolism, proliferation, cell survival and angiogenesis [8].AIM2 also promotes macrophage recruitment during experimental head and neck squamous cell carcinoma [9]. Additional inflammasome-independent roles for AIM2 have been identified during T cell differentiation, although these studies have reported conflicting outcomes. For example, AIM2 was implicated in promoting Treg stability through diminishing Akt-mTOR signaling and modulating metabolism during experimental autoimmune encephalomyelitis and T-cell mediated colitis [10, 11]. Another study argued AIM2 suppresses Treg differentiation during T-mediated colitis, also through modulating metabolism [10, 11]. AIM2 was recently found to promote Th17 differentiation of naïve CD4^+^ cells through the direct regulation of RORγt, the lineage defining transcription factor for Th17 cells [12]. AIM2 was also implicated in driving Tfh differentiation during systemic lupus erythematosus [13]. Thus, AIM2 performs important functions during T cell differentiation through regulating metabolism and the expression of master transcription factors, but this may depend on the strength of TCR stimulus during activation (e.g., antibody-mediated vs. peptide-MHC) and the environmental context (e.g. presence of specific cytokines and DAMPs) by which the T cell is activated. Importantly, several of the experiments supporting a role for AIM2 in T cells were performed using non-antigen-specific in vitro assays, and thus it is unclear how AIM2 impacts T cell functions during defined antigen-specific activation.

In this study, we explored the intrinsic role of AIM2 in CD4^+^ T cells during antigen-specific activation using *Aim2^-/-^* mice crossed to OT-II mice, which only contain CD4^+^ T cells with OVA-specific TCRs. At steady state, we observed that *Aim2^-/-^*/OT-II displayed no alterations in single positive (CD4^+^) to double positive (CD4^+^ CD8α^+^) thymocyte ratios compared to OT-II mice. However, naïve *Aim2^-/-^*/OT-II mice had decreased numbers of CD4^+^ T cells in the spleen and a decreased number of CD24^hi^/Qa-2^lo^ recent thymic emigrants to the spleen. Compared to OT-II CD4^+^ controls, *Aim2^-/-^*/OT-II CD4^+^ T cells co-cultured with OVA-incubated dendritic cells in vitro produced less IL-2 and displayed decreased expression of the high affinity IL-2 receptor *Il2ra*. However, we failed to detect differences in proliferation or differentiation of *Aim2^-/-^*/OT-II and OT-II CD4^+^ T cell adoptively transferred into C57Bl/6J mice immunized with CFA OVA peptide. Lastly, *Aim2^-/-^*/OT-II T cells controlled the growth of implanted OVA-NOOC2 tumor cells to a similar capacity as WT OT-II cells. Altogether, our results indicate that AIM2 plays a role in promoting CD4^+^ T cell thymic selection, retention in the spleen and antigen-specific activation in vitro. However, these functions may not impact CD4^+^ T cell proliferation and differentiation during primary immunization or anti-tumor control in vivo.

## Materials and Methods

### Mice

All animal protocols were approved by the University of Arizona Institutional Animal Care and Use Committee (IACUC) in accordance with the US National Institutes of Health Guide for Care and Use of Laboratory Animals. OT-II TCR tg mice were generated on the C57BL/6J CD45.1 Rag1KO background as previously described [14]. Briefly, the hCD2-OT-IIα TCR and hCD2-OT-IIβ constructs were in injected into C57BL/6J embryos. Founder mice were then crossed to C57BL/6J Rag1KO mice and then crossed to C57BL/6J CD45.1 Rag1KO mice. To generate *Aim2^-/-^* OT-II mice, *Aim2^-/-^* mice on the C57BL/6J background were crossed with the OT-II mice.

#### Cell lines and culture

The NOOC2-OVA cell line was generated at the University of Texas M.D. Anderson Cancer Center, Houston TX. Briefly, the parental NOOC2 cells were generated from macroscopic oral squamous cell carcinoma lesions excised from C57BL/6J mice given the oral carcinogen 4NQO in drinking water (50 µg/mL) for 16 weeks. The oral lesions were digested into single cell suspensions, and single cell clones were grown in culture and screened for their ability to produce tumors when implanted in syngeneic C57BL/6J hosts [15, 16]. NOOC2-OVA were generated using stable retroviral transduction of OVA plasmid. OVA protein expression was verified by western blot (Data not shown). NOOC2-OVA cells were maintained in 60% IMDM (Corning) and 30% F12 nutrient mix (Corning) with 5% FBS, supplemented with 4 µg/mL puromycin, 5 µg/mL insulin, 40 ng/mL hydrocortisone, 5 ng/mL EGF and 100 U/mL penicillin-streptomycin.

#### Adoptive transfers, peptides, CFA and immunizations

Single cell suspensions were generated and pooled from the spleens, brachial, axillary, and inguinal lymph nodes of OT-II mice. CD4^+^ T cells were then enriched using CD4 T cell isolation kit (Miltenyi Biotec) and MACS separation columns (Miltenyi Biotec). CD4^+^ T cells were stained with Tag-it Violet (Biolegend) (5 µM) according to the manufacturer’s instructions. A total 1 x 10^5^ Tag-it Violet (Biolegend) labelled CD4^+^ T cells were retro-orbitally transferred into recipient C57BL/6J mice. One day later, mice were immunized with 20 µg of OVA_323-339_ peptide (ISQAVHAAHAEINEAGR, purchased from InvivoGen, San Diego, CA) in Complete Freund’s adjuvant (CFA) (Sigma-Aldrich, St. Louis, MO) on the back flank. At 60 h (proliferation analysis) or 6 d (differentiation analysis) postimmunization, the draining lymph nodes (brachial, axillary, inguinal) were harvested, and CD4^+^ T cells were enriched and stained with antibodies for analysis by flow cytometry.

#### In vivo proliferation analysis

CD4^+^ T cells were labelled with Tag-it Violet and analyzed for proliferation by measuring the subsequent dilution of the fluorescent dye by flow cytometry. The total number of daughter and undivided cells, the number of responding T cells (number of original T cells that divided due to stimulus), and the proliferative capacity (the average number of daughter cells generated per responder) were calculated as previously described [17].

#### Bone marrow derived dendritic cell generation

We generated bone marrow derived dendritic cells as previously described [18]. Briefly, bone marrow was isolated from the femurs and tibia of C57BL/6J mice and processed to obtain single cell suspensions. A total of 4 x 10^6^ bone marrow cells were plated on a 15 cm plate in RPMI with 10% FBS and 1 % Penicillin/Streptomycin, 50μM β-mercaptoethanol and 20 ng/mL of murine GM-CSF (Peprotech, 315-03) for 10 days with new media being added at day 3, day 6 and day 8. Differentiated bone marrow derived dendritic cells were used for CD4^+^ T cell coculture assays.

#### IL-2 ELISA

CD4^+^ T cells (5 x 10^4^) were enriched from OT-II or *Aim2^-/-^*OT-II mice and co-cultured with 1 x 10^5^ bone marrow derived dendritic cells generated from C57BL/6J mice. Cocultures were performed in triplicate in a 96-well round bottom plate containing RPMI 1640 with 5% FBS, penicillin/streptomycin (product), 50 mM 2-ME (Fisher Scientific) in the presence of differing amounts of Ovalbumin protein (0 µM, 1 µM, 5 µM, 10 µM, 15 µM, and 20 µM). The supernatants were collected and analyzed for IL-2 cytokine production by ELISA (R&D Systems) after 16-18 hours of coculture at 37°C. The ELISA was performed according to the manufacturer’s recommendations in duplicate. ELISA plates were analyzed by spectrometer Biotek Synergy H1 plate reader according to the manufacturer’s instructions.

#### RNA Isolation

Total RNA was isolated from CD4^+^ T cell and bone marrow derived dendritic cell cocultures after supernatant harvest at 16-18 hours of coculture with OVA_323-339_ coculture. RNeasy Plus Mini Kit (Qiagen, 74136) was utilized according to the manufacturer’s protocol for isolation from cells. RNA concentrations were determined using a spectrophotometer. cDNA was synthesized at a 200 µg/µl template using the iScript Reverse Transcription Supermix for RT-qPCR (Bio-Rad, 1706840) and diluted 1:10 with molecular grade water before performing TaqMan RT-qPCR.

#### Flow Cytometry

Single-cell suspensions were generated from the thymus and spleens of OT-II or *Aim2^-/-^* OT-II mice. Single-cell suspensions were stained using Zombie Violet (Biolegend) for 30 min at 4°C. Cells were then fixed using 4% paraformaldehyde for 15 min at 4°C. Cells were then washed 2 times and stained for 30 min at 4°C with corresponding antibodies. Antibodies used for this study included anti-CD4 (RM4-5), CD8α (53-6.7), CCR7/CD197 (4B12), TCR Vα2 (B20.1), TCR Vβ5 (MR9-4), PD-1, Qa-2, CD44, CD45.1, CD45.2 and CD185/CXCR5 (all purchased from Biolegend). Staining for CCR7/CD197 was performed prior to Zombie Violet staining, fixing, and other surface staining at 37°C for 30 min. After surface staining, cells were washed before being stored at 4°C. Flow cytometry was performed on a Cytek Aurora (Cytek) instrument, and data was analyzed with FlowJo v11 (Becton Dickinson).

#### Tumor Injection

A total of 1.8 x 10^6^ NOOC2-OVA cells were injected into the right flanks of C57BL/6J recipient mice. Tumors were allowed to establish for 8 days. Then after 8 days, enriched CD4^+^ OT-II cells, *Aim2^-/-^*OT-II cells, or PBS vehicle were retro-orbitally injected into the mice. The tumors were then allowed to grow for 56 days. Tumors were measured using calipers 3 times a week every other day for the duration of the experiment. Mice were euthanized at the endpoint or when moribund for tumor size measurements.

#### Quantitative Real-Time PCR

qPCR analysis was performed on cDNA from cell coculture samples using the TaqMan expression assays (Applied Biosystems). The primer targets used in this study included *Il2ra* and *Prdm1*. Reactions were run in technical duplicates on a Quantstudio 3 System in a 96-well 0.2 ml block. Relative expression was calculated with the ΔΔCT method using *Gapdh* expression as the endogenous control. Relative expression was plotted as the mean +/- SEM.

## Results

### AIM2 does not impact thymocyte development

To evaluate the impact of AIM2 on antigen-specific T cell activation, we crossed *Aim2^-/-^* mice with OT-II/RAG transgenic mice, which contain CD4^+^ T cells that exclusively express an ovalbumin (OVA)-specific TCR that recognizes the OVA_323-339_ peptide fragment (I-A^b^ restricted) and no other lymphocytes due to RAG deficiency. This resulted in the generation of mice that lack AIM2 and express the OT-II TCR in CD4^+^ T cells (*Aim2^-/-^*/OT-II). T cells undergo development in the thymus starting as double negative CD4/CD8, then becoming double positive for CD4/CD8 before finally transitioning to a single positive CD4/CD8 stage after selection. OT-II mice do not generate single positive CD8 T cells due to the nature of their genetic background [14]. Thus, we characterized the resulting *Aim2^-/-^*/OT-II by first measuring thymocyte development. To do this, we isolated the thymus of 6–8-week-old WT OT-II and *Aim2^-/-^*/OT-II mice and measured CD4 and CD8 expression on thymocytes by flow cytometry (**Fig. 1A**). There were no significant differences in the total number of thymocytes between the *Aim2^-/-^*/OT-II and WT OT-II mice (**Fig. 1B**) nor were there any differences in the number of CD4/CD8 double positive or CD4^+^ single positive cell numbers between these mice (**Fig. 1C-D**). These results suggest that AIM2 does not significantly impact thymocyte development.

**Figure 1.**
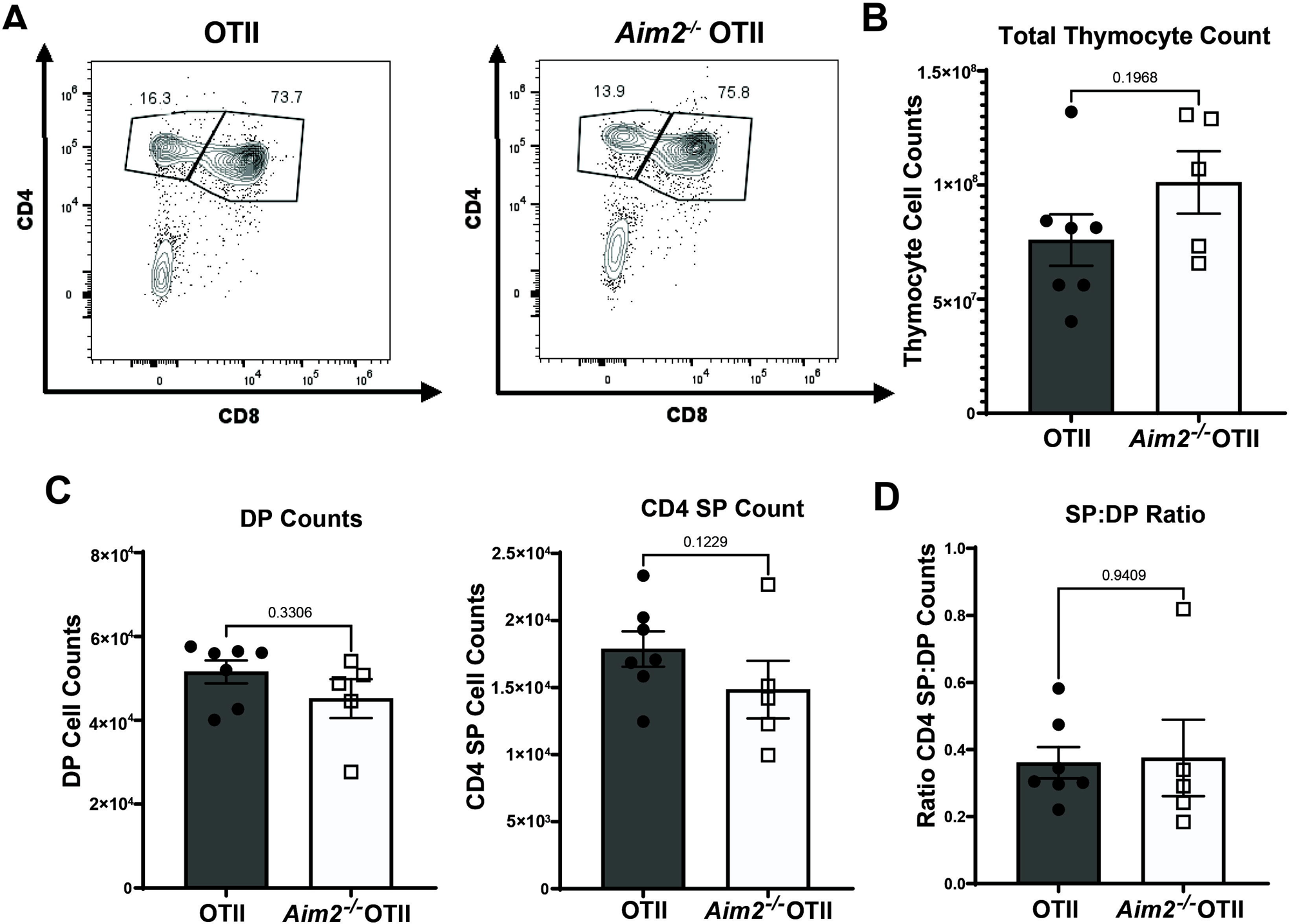
*Aim2^-/-^* /OT-II display no differences in thymocyte development. (A) Representative CD8a versus CD4 flow cytometry plots are shown for either WT OT-II (left) or *Aim2^-/-^* OT-II (right) thymocytes after gating on lymphocytes using side scatter vs. forward scatter (FSC), FSC-height vs FSC-area doublet discriminator and Live/Dead Violet (data not shown). Percentages are inset for each population. (B) The absolute numbers of thymocytes shown. (C) DP (left) and CD4 SP (right) thymocytes counts are shown. (D) The ration of CD4^+^ SP counts to CD4^+^CD8a^+^ DP counts are shown. Bars represent mean values ± SEM. Each dot represents a single mouse 6-7 weeks of age (OT-II, n= 7; *Aim2^-/-^* OT-II, n= 5). Analyzed by unpaired *t* test.

### AIM2 regulates positive selection of thymocytes

To further characterize thymic development, we assessed thymic selection of single positive CD4^+^ T cells using gating strategies based on publications in the field [14]. Briefly, we gated on CD4^+^ CD5^hi^ thymocytes to analyze both SPs and DPs that have had signaling through their TCR (**Fig. 2A**). We then gated on TCR^hi^ CCR7^+^, which restricts the analysis to thymocytes that are experiencing or have undergone positive selection (**Fig. 2A**). Lastly, we used CD69 expression to distinguish thymocytes actively undergoing positive selection (**Fig. 2A**). Upon evaluation of the CD4^+^ CD5^hi^ TCRvβ5^hi^ CCR7^+^ CD69^+^ population, we found that a greater percentage of *Aim2^-/-^*/OT-II thymocytes were auditioning for selection, however their overall numbers were similar to WT OT-II thymocytes (**Fig. 2B-C**). When assessing the positively selected CD4^+^ CD5^hi^ TCR^hi^ CCR7^+^ CD69^-^ population, we found that *Aim2^-/-^*/OT-II thymocytes had a decreased number and percentage of SP CD4^hi^ cells compared to WT OT-II thymocytes (**Fig. 2B,D**). These findings suggest there were fewer positively selected CD4^+^ thymocytes in *Aim2^-/-^*/OT-II mice compared to WT OT-II mice. Upon analyzing CD5 expression levels (MFI) among these CD69^+^ and CD69^-^ CD4^+^ thymocytes as a measurement of TCR signal strength in response to selection. We found no significant differences in CD5 expression levels between thymic WT OT-II and *Aim2^-/-^*/OT-II CD4^+^ thymocytes (**Fig. 2E**). This suggests that while AIM2 may promote positive selection in the thymus, this cannot be explained by alterations in TCR signal strength. Overall, these results indicate that while AIM2 does not impact the total numbers of thymocytes, yet it does promote positive selection of SP CD4^+^ thymocytes.

**Figure 2.**
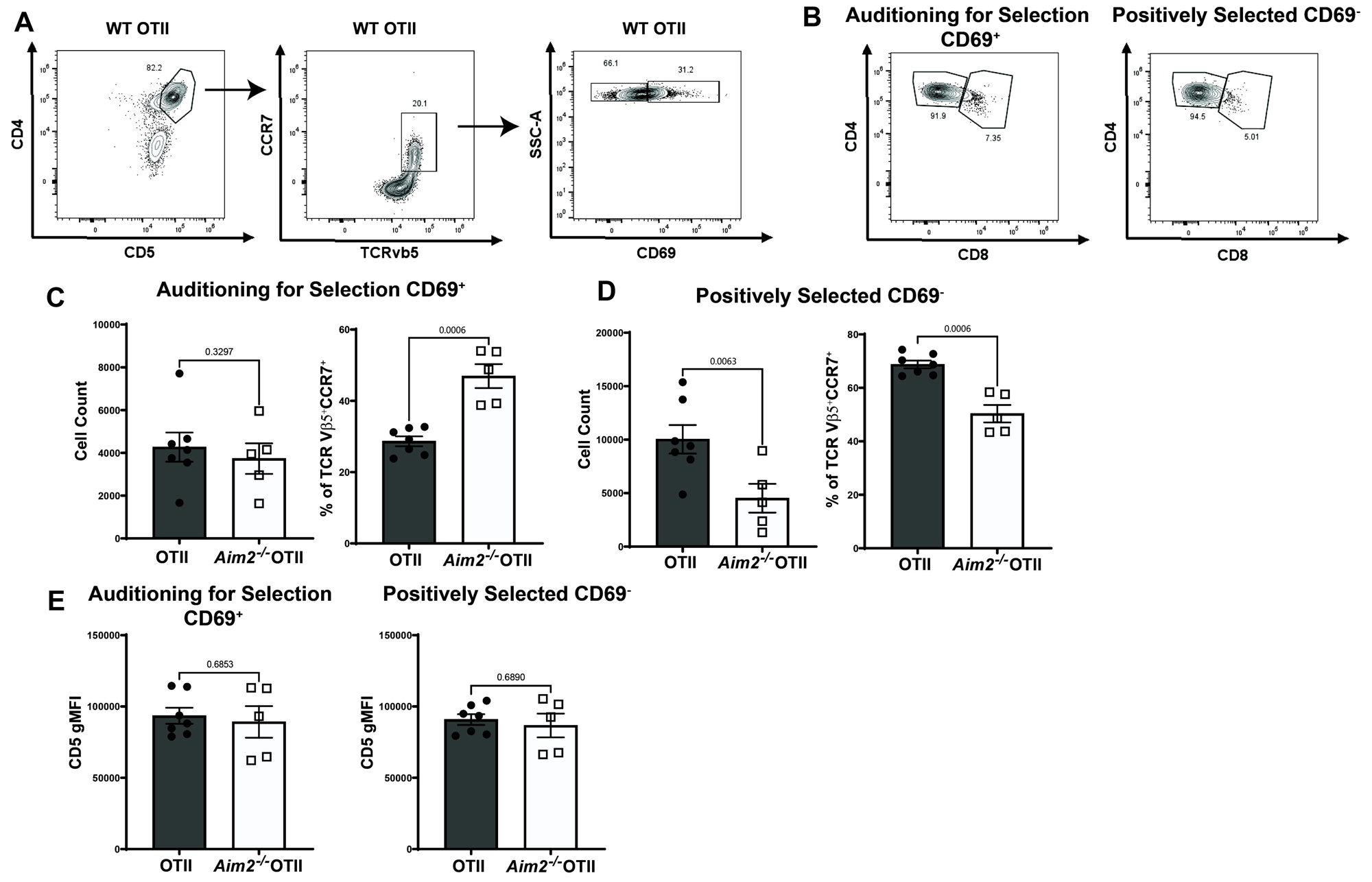
*Aim2^-/-^*OT-II mice display reduced populations of positively selected CD4 thymocytes and increased numbers of thymocytes auditioning for selection. (A) Representative gating strategy of OT-II thymocytes (WT shown as example), after gating on lymphocytes using side scatter vs. forward scatter (FSC), FSC-height vs. FSC-area doublet discriminator, and Live/Dead violet. CD4^+^ CD5^hi^ WT thymocytes (left) were gated for TCR and CCR7 expression (center), then gated on CD69 expression (right). (B) CD8 vs. CD4 expression is shown for CD4^+^ CD5^hi^ TCRvβ5^hi^ CCR7^+^ CD69^+^ (left) and CD69^-^ (right) subsets. (C) Total cell counts (left) and % of parental populations (right) are shown for CD4^+^ CD5^hi^ TCRvβ5^hi^ CCR7^+^ CD69^+^ gated WT and *Aim2^-/-^* OT-II thymocyte subsets. (D) Total cell counts (left) and % of parental populations (right) are shown for CD4^+^ CD5^hi^ TCRvβ5^hi^ CCR7^+^ CD69^-^ gated WT and *Aim2^-/-^* OT-II thymocyte subsets. (E) CD5 gMFI for CD4^+^ CD5^hi^ TCRvβ5^hi^ CCR7^+^ CD69^+^ (left, Auditioning for Selection) or CD69^-^ (right, Positively Selected) gated WT and *Aim2^-/-^* OT-II thymocyte subsets. Each dot represents a single mouse 6-7 weeks of age (OT-II, n=6-9; *Aim2^-/-^*OT-II, n=7), with bars representing mean ± SEM. Analyzed by unpaired *t* test.

### AIM2 promotes the emigration of CD4^+^ thymocytes to the spleen

After thymic development, CD4^+^ T cells migrate to secondary lymphoid organs including the lymph nodes and spleen. Therefore, we next measured splenocyte numbers and recent thymic emigrants to the spleen. While we found a trending reduction, but no significant difference in the total number of splenocytes in *Aim2^-/-^*/OT-II mice (**Fig. 3A**). There was a significant decrease in the number of *Aim2^-/-^*/OT-II CD4^+^ T cells compared to WT OT-II mice (**Fig. 3B**). Recent thymic emigrants in the spleen can be identified by low expression of (Qa-2^lo^) and high expression of (CD24^hi^) (**Fig. 3C**) [19]. Because we detected reduced splenic CD4^+^ T cells in the absence of AIM2, we next assessed the recent thymic emigrant population and found that *Aim2^-/-^*/OT-II spleens had significantly lower percentages and fewer numbers of Qa-2^lo^ CD24^hi^ recent thymic emigrants compared to WT OT-II mice (**Fig. 3D**). Conversely, *Aim2^-/-^*/OT-II spleens also had a significantly greater percentage, but equal numbers of Qa-2^hi^ CD24^lo^ cells, which represent more mature thymic emigrants (**Fig. 3D**). Because tonic TCR signaling plays an important role in homeostatic survival of T cells within the spleen, we next assessed CD5 expression levels as an indicator of tonic TCR signaling in these CD4^+^ populations [20]. Similar to the thymus, we found no significant difference in CD5 expression levels in splenic CD4^+^ T cells from WT OT-II and *Aim2^-/-^*/OT-II mice (**Fig. 3E left**). We then confirmed this using a CD5 versus Ly6C gating strategy as this analysis allows for a broader detection range of tonic TCR signaling compared to CD5 alone (i.e., high tonic signaling) [14, 21]. This analysis revealed that there was no significant difference in the CD5^hi^ Ly6C^lo^ CD4^+^ T cell population experiencing high tonic signaling among the WT OT-II and *Aim2^-/-^* OT-II spleens suggesting that tonic signaling is not altered within *Aim2^-/-^*OT-II T cells (**Fig. 3E**). Overall, these results indicate that *Aim2^-/-^*/OT-II cells have reduced thymic emigration, but the CD4^+^ T cells that arrive at the spleen are not only capable, but potentially hastened in their maturation as evidenced by their increased expression of Qa-2. Thus, the reduced number of CD4^+^ T cells observed in the *Aim2^-/-^*/OT-II spleens may be due to the reduction in thymic selection of SP CD4^+^ and subsequent reduction in recent thymic emigrants to the spleen.

**Figure 3.**
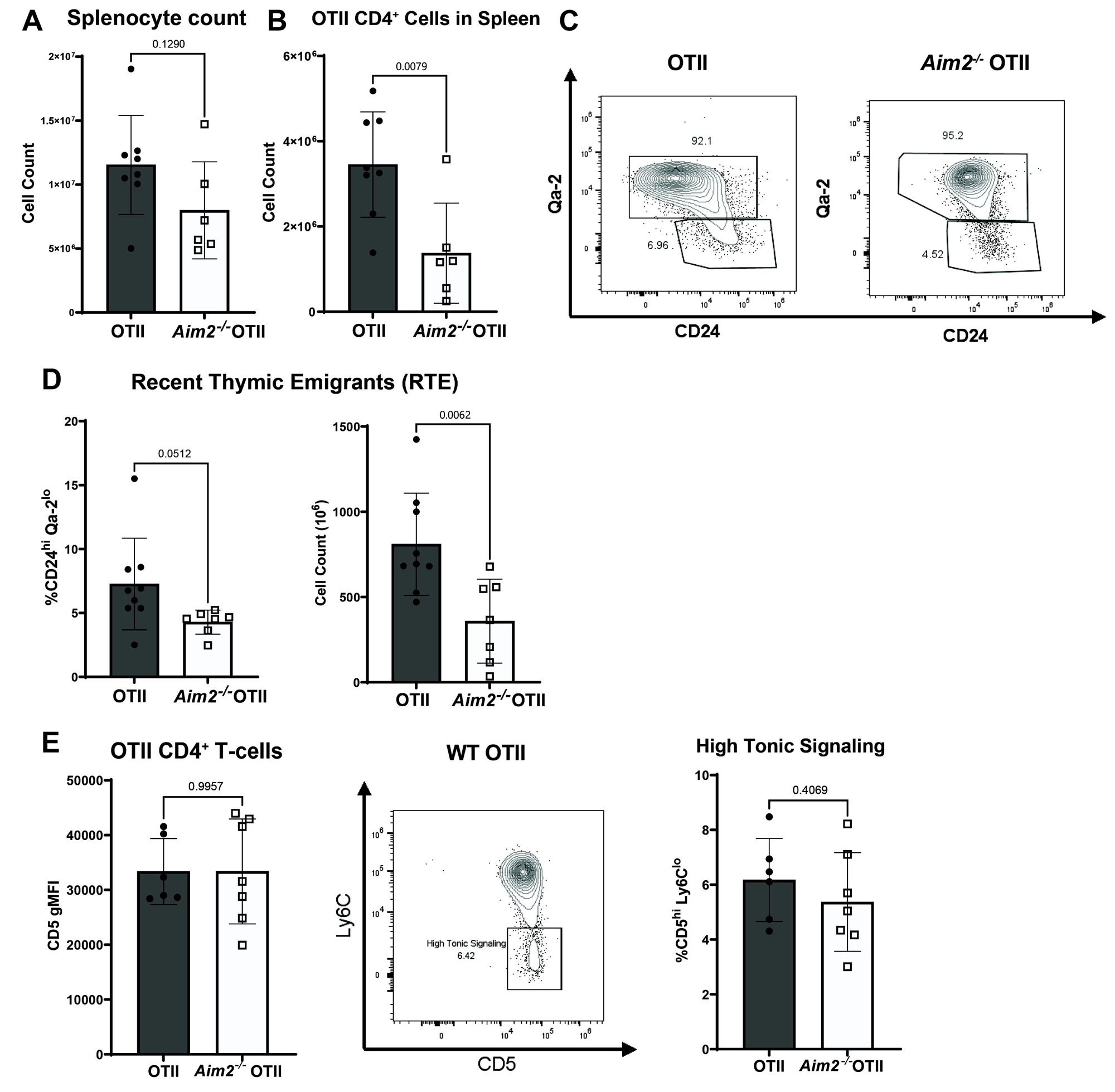
*Aim2^-/-^* OT-II mice have fewer CD4^+^ cells in the spleen and fewer recent thymic emigrants. (A) Total number of splenocytes counted from spleens of OT-II (n=8) and *Aim2^-/-^* OT-II (n=6). (C) Representative CD24 vs. Qa-2 flow cytometry plots are shown for WT OT-II (left) and *Aim2^-/-^* OT-II (right) splenocytes after gating on lymphocytes using side scatter vs. forward scatter (FSC), FSC-height vs. FSC-area doublet discriminator, Live/Dead violet and TCRvβ5 vs. CD4 (data not shown). (D) Frequency of peripheral CD4^+^ T cells that are recent thymic emigrants (RTEs) (left) and their total numbers (right). (E) CD5 gMFI for CD4^+^ TCRvβ5^+^ OT-II T-cells (WT and *Aim2^-/-^*OT-II) from the spleen are shown (left). CD5 vs Ly6C expression is shown after gating on CD5^hi^ Ly6C^-^ CD4^+^ T cells experiencing high tonic signaling (middle) after gating on SSC vs. FSC, FSC-H vs. FSC-A doublet discriminator, Live/Dead violet, and TCRvβ5 vs. CD4. Representative WT OTII CD4^+^ T cells are shown. The frequency of CD4^+^ T cells that are CD5^hi^ Ly6C^-^ are shown (right). Each dot represents a single mouse 6-7 weeks of age (OT-II, n=6-9; *Aim2^-/-^* OT-II, n=7), with bars representing mean ± SEM. Data was analyzed by unpaired *t* test.

### AIM2 promotes IL-2 production from CD4^+^ T cells in response to antigen presentation in vitro

We asked if AIM2 has an intrinsic impact on antigen-specific CD4^+^ T cell activation in vitro. This was accomplished by co-culturing WT OT-II or *Aim2^-/-^*/OT-II CD4^+^ T cells with bone marrow derived dendritic cells incubated with 0-20 µM whole OVA for 16 hours. We then evaluated CD4^+^ T cell activation by measuring IL-2 production in the supernatant by ELISA. This timepoint was chosen because IL-2 production from CD4^+^ T cells often peaks around 6-16 hours after antigen challenge and is undetectable or significantly decreased by 24 hours due to consumption [22]. Across the OVA titration range, WT OT-II CD4^+^ T cells produced more IL-2 than the *Aim2^-/-^*/OT-II CD4^+^ T cells at 16 hours post activation (**Fig. 4A**). Indeed, IL-2 production in the *Aim2^-/-^*/OT-II CD4^+^ T cells was reduced in response magnitude compared to WT, as quantified by the area under the curve for the overall response to the peptide range (**Fig. 4B**). We also harvested total RNA from these cocultures and measured the mRNA levels of IL2Rα, using qPCR, which is an important component of the high affinity IL-2 receptor that is upregulated upon TCR activation Expression of *Il2ra* RNA was reduced in *Aim2^-/-^*/OT-II CD4^+^ T cells compared to WT cells suggesting *Aim2^-/-^*/OT-II CD4^+^ T cells express less of the high affinity IL2 receptor following activation (**Fig. 4C**).

**Figure 4.**
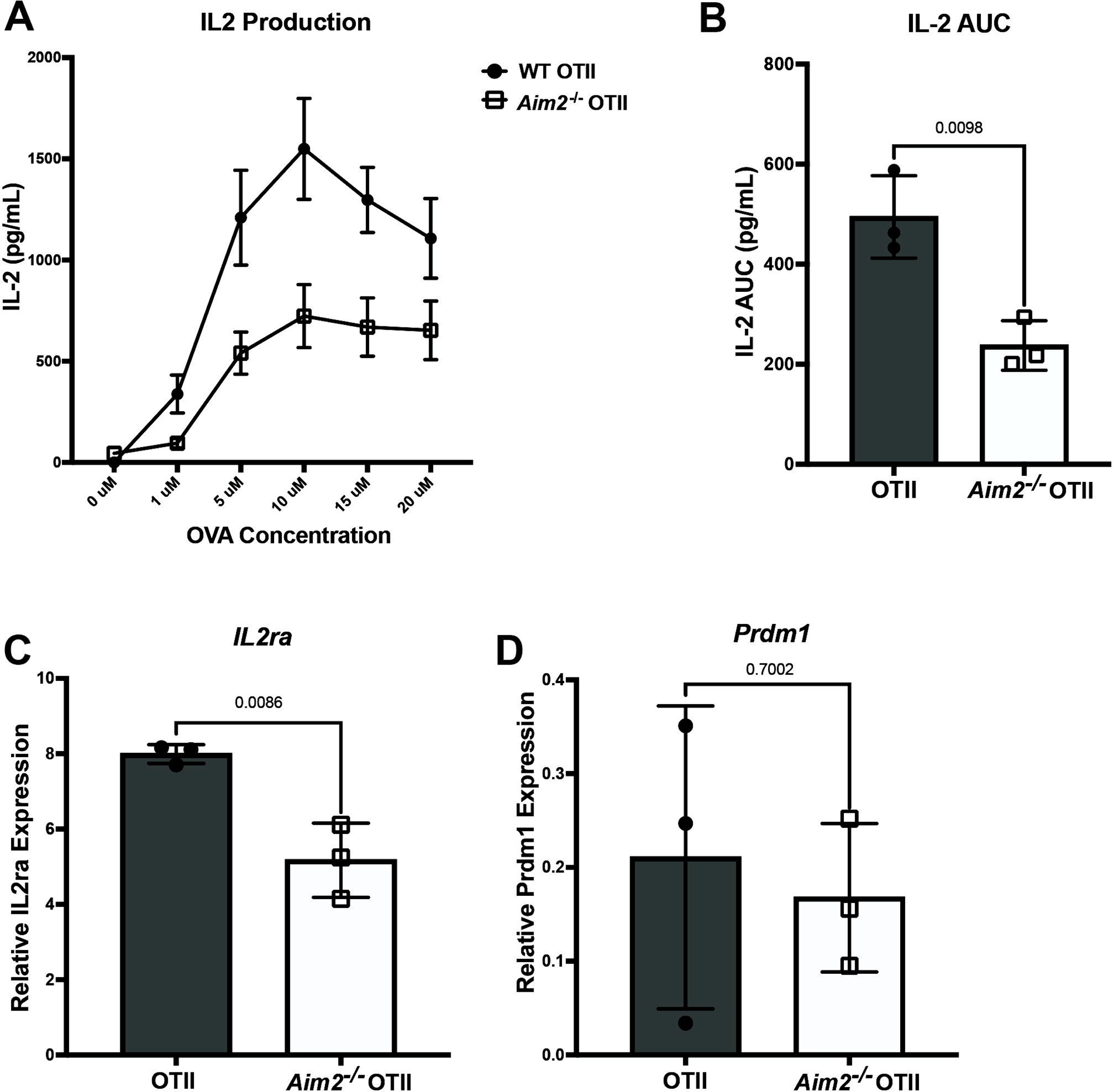
*Aim2^-/-^* OT-II CD4 T cells produce less IL-2 and express reduced *Il2ra* in response to antigen presentation in vitro. (**A**) Representative curve for IL-2 production by OT-II (n=3) and *Aim2^-/-^* OT-II (n=3) CD4 T cells in response to indicated concentrations of ovalbumin (OVA) presented by bone marrow derived dendritic cells from C57Bl/6J mice. (**B**) Area under the curve (AUC) analysis for the OVA dose response shown as a the measurement of the response magnitude. (**C**) RT-qPCR analysis of *Il2ra* mRNA expression from OT-II or *Aim2^-/-^* OT-II CD4 T cells activated as in (A). (**D**) RT-qPCR analysis of *Prdm1* mRNA expression from OT-II or *Aim2^-/-^* OT-II CD4 T cells activated as in (A). Each dot represents one mouse. Data are presented as means +/-SEM and analyzed by unpaired *t* test.

IL-2 expression is very transient due to being consumed and its ability to induce a potent negative feedback loop, which downregulates its own expression [22]. This negative feedback loop occurs in response to IL-2 binding to its high affinity IL-2R, which results in the activation of signal transducer and activator of transcription-5 (STAT5) and induction of B lymphocyte maturation protein-1 (BLIMP-1, *Prdm1*). BLIMP-1 then represses IL-2 expression [22]. AIM2 regulates B cell differentiation by modulating Bcl-6-Blimp-1 and is highly expressed in B cells in lupus patients [23]. To determine if the decreased IL-2 and *Il2ra* expression in the *Aim2^-/-^*/OT-II CD4^+^ T cells was associated with enhancement of this negative feedback loop through BLIMP-1, we measured the expression of *Prdm1* in WT OT-II and *Aim2^-/-^*/OT-II CD4^+^ T cells.

We found no difference in the expression of *Prdm1* between WT or *Aim2^-/-^* OT-II CD4^+^ T cells (**Fig. 4D**) suggesting that the decrease in *Il2ra* was not due to greater activation of the BLIMP-1-dependent negative feedback loop. Taken together, *Aim2^-/-^*/OT-II CD4^+^ T cells have a defect in IL-2 production and reduced *IL2ra* expression in response to antigen-specific activation in vitro. These findings suggest AIM2 intrinsically promotes IL-2 production or signaling in CD4^+^ T cells, which could promote T cell function.

### AIM2 does not impact CD4^+^ T cell proliferation or differentiation in response to CFA/OVA immunization

The primary role of IL-2 is to promote T cell proliferation and differentiation during the early stages of T cell activation. Because we found reduced IL-2 production in *Aim2^-/-^*/OT-II CD4^+^ T cells in vitro, we asked if this translated into a defect in the proliferative capacity of these cells in response to immunization with OVA peptide in vivo. Specifically, we adoptively transferred Tag-It Violet-labeled WT or *Aim2^-/-^*/OT-II CD4^+^ T cells into C57BL/6J recipients, immunized the recipient mice with OVA_323-339_ in Complete Freund’s Adjuvant (CFA) and quantified OT-II proliferation by Tag-It violet dye dilution at 60 h postimmunization (**Fig. 5A**). We enumerated the daughter cells, which displayed diluted Tag-it dye, TCRVβ5 and the congenic marker CD45.1 and back-calculated the number of cells that would have had to respond to generate that number of daughter cells. We also calculated the average number of daughter cells that came from each responder [14, 17]. We found a trending, but non-significant (p=0.0576) increase in the number of daughter cells generated by *Aim2^-/-^*/OT-II CD4^+^ T cells. No significant differences were detected between the number of responders or the proliferative capacity among the WT and *Aim2^-/-^*/OT-II CD4^+^ T cells, overall suggesting the reduction in IL-2 production observed in *Aim2^-/-^*/OT-II cells in vitro does not result in their defective proliferation in vivo (**Fig. 5A**). However, the *Aim2^-/-^*/OT-II cells displayed a significantly greater number of undivided cells compared to WT OT-II controls. This overall suggests that compared to WT OT-II controls, fewer *Aim2^-/-^*/OT-II cells became activated, but those that were activated underwent robust proliferation. This is supported by our finding that *Aim2^-/-^*/OT-II produced the same number of daughters as the WT OT-II cells. Alternatively, these results could reflect an enhancement of survival in the transferred *Aim2^-/-^*/OT-II population before immunization.

**Figure 5.**
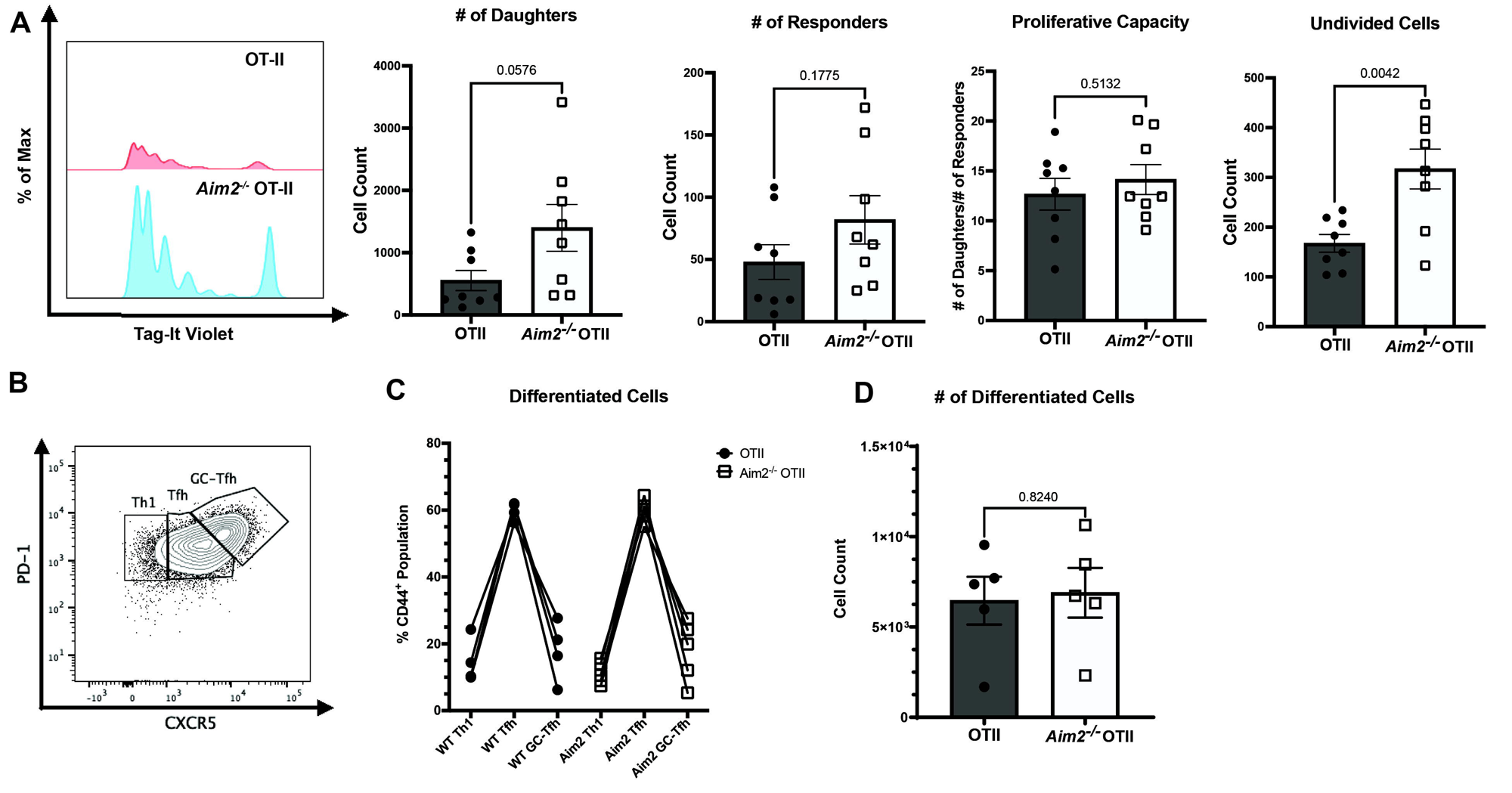
*Aim2^-/-^*OT-II CD4 T cells undergo normal proliferation and differentiation in response to immunization in vivo. (**A**) Left to right, Concatenated in vivo proliferation histogram plot of WT OT-II and *Aim2^-/-^*OT-II CD4 T cells adoptively transferred into C57BL/6J recipient mice prior to immunization with OVA_323-339_ peptide in CFA, as measured by Tag-It Violet dye dilution. The number of divided cells, the calculated number of responders that would generate the daughter cell population (see *Materials and Methods*), the average number of daughter cells per responder (proliferative capacity, see *Materials and Methods*), and the number of undivided cells are shown. Tag-it Violet cells were identified by gating on forward scatter (FSC) vs. side scatter, FSC-height vs. FSC-area doublet discriminator, Live/Dead, CD4^+^CD8^-^, Vα2^+^Vβ5^+^, and CD45.1^+^. Each dot represents a recipient mouse (n=8). (**B**) Representative gating of adoptively transferred WT OT-II or *Aim2^-/-^*OT-II CD4 T cells 6 days after OVA-CFA immunization of C57BL/6J recipient mice for Th1(CXCR5^lo^ PD-1^lo^), Tfh (CXCR5^Int^ PD-1^Int^), and germinal center-Tfh (GC-Tfh) (CXCR5^hi^ PD-1^hi^) populations after pregating on FSC vs. side scatter, FSC-height vs. FSC-area doublet discriminator, Live/Dead, CD4^+^CD8^-^, Vα2^+^Vβ5^+^, and CD44^hi^. (**C**) The percentage of adoptively transferred populations that differentiated into Th1, Tfh, and GC-Tfh are shown. (**D**) The absolute number of adoptively transferred cells enumerated on day 6. Each dot represents one mouse (n=5 mice/group). Data are presented as means +/-SEM and analyzed by unpaired *t* test.

IL-2 directs the differentiation of CD4^+^ T cells [22, 24, 25]. Other studies using distinct disease models and non-antigen specific in vitro skewing assays have implicated differing impacts of AIM2 on CD4^+^ differentiation [10–13]. Thus, we next assessed if the in vitro regulation of IL-2 by AIM2 translated into alterations in CD4^+^ differentiation in response to Ag-specific activation in vivo. To do this, we adoptively transferred WT OT-II or *Aim2^-/-^*/OT-II CD4^+^ T cells into C57BL/6J recipients and immunized the recipient mice with OVA_323-339_ in CFA the following day. At 6 days postimmunization, we isolated the draining lymph nodes and assessed CD4^+^ T cell differentiation into Th1 and Tfh cells as previously reported using OT-II mice [14, 26]. We identified the adoptively transferred cells through their clonotypic TCR Vα2 and Vβ5 expression as well as the congenic CD45.1 marker. We also utilized CD44^hi^ staining to observe cells that responded to stimuli and underwent differentiation. Lastly, we used CXCR5 and PD-1 expressions to identify Th1(CXCR5^lo^ PD-1^lo^), Tfh (CXCR5^Int^ PD-1^Int^), and germinal center-Tfh (GC-Tfh) (CXCR5^hi^ PD-1^hi^) CD4^+^ T cells. We found no significant differences in the percentages and numbers of Th1, Tfh or GC-Tfh populations in the transferred WT OT-II and *Aim2^-/-^*/OT-II CD4^+^ cells, indicating AIM2 had no measurable impact on CD4^+^ T cell differentiation in response to OVA immunization in vivo (**Fig. 5B-D**).

### AIM2 does not impact antigen-specific effector functions of CD4^+^ T cells during syngeneic tumor challenge

Since we observed no differences in OVA immunization-induced proliferation or differentiation between WT and *Aim2^-/-^*/OT-II cells, we then asked if AIM2 impacted antigen-specific effector functions against other immunological insults. CD4^+^ T cells are key mediators for anti-tumor immunity through their ability to guide other immune effectors, including cytotoxic CD8 T cells [27]. To address this, we utilized a syngeneic tumor injection model in which we subcutaneously injected OVA-expressing murine head and neck tumor cell line NOOC2 (NOOC2-OVA) cells into the back flank of C57BL/6J mice. Once the tumor cells were established at 10 days, we adoptively transferred WT OT-II or *Aim2^-/-^*/OT-II cells into the tumor bearing mice via intravenous injection and measured tumor growth every other day until the tumors reached sufficient size that required the mice to be euthanized. Additional tumor-bearing mice were intravenously injected with PBS (no OT-II cells) as a control. While we saw a significant reduction in NOOC2-OVA tumor growth in animals adoptively transferred with either WT or *Aim2^-/-^*/OT-II cells compared to PBS-injected control mice, there was no significant difference in tumor growth in mice given WT OT-II or *Aim2^-/-^*/OT-II cells (**Fig. 6A-B**). While no survival differences were detected between these mice, we had to end the study at 56 days post-tumor injection due to the required humane endpoint determined by our IACUC protocol. This suggests that despite the reduction in positively selected CD4^+^ T cells in the thymus, reduced CD4^+^ T cells in the spleen and a dramatic reduction in IL-2 production by *Aim2^-/-^*/OT-II cells in vitro, these defects did not translate into measurable differences in proliferation, differentiation or antigen-specific anti-tumor immunity against NOOC2 challenge in vivo.

**Figure 6.**
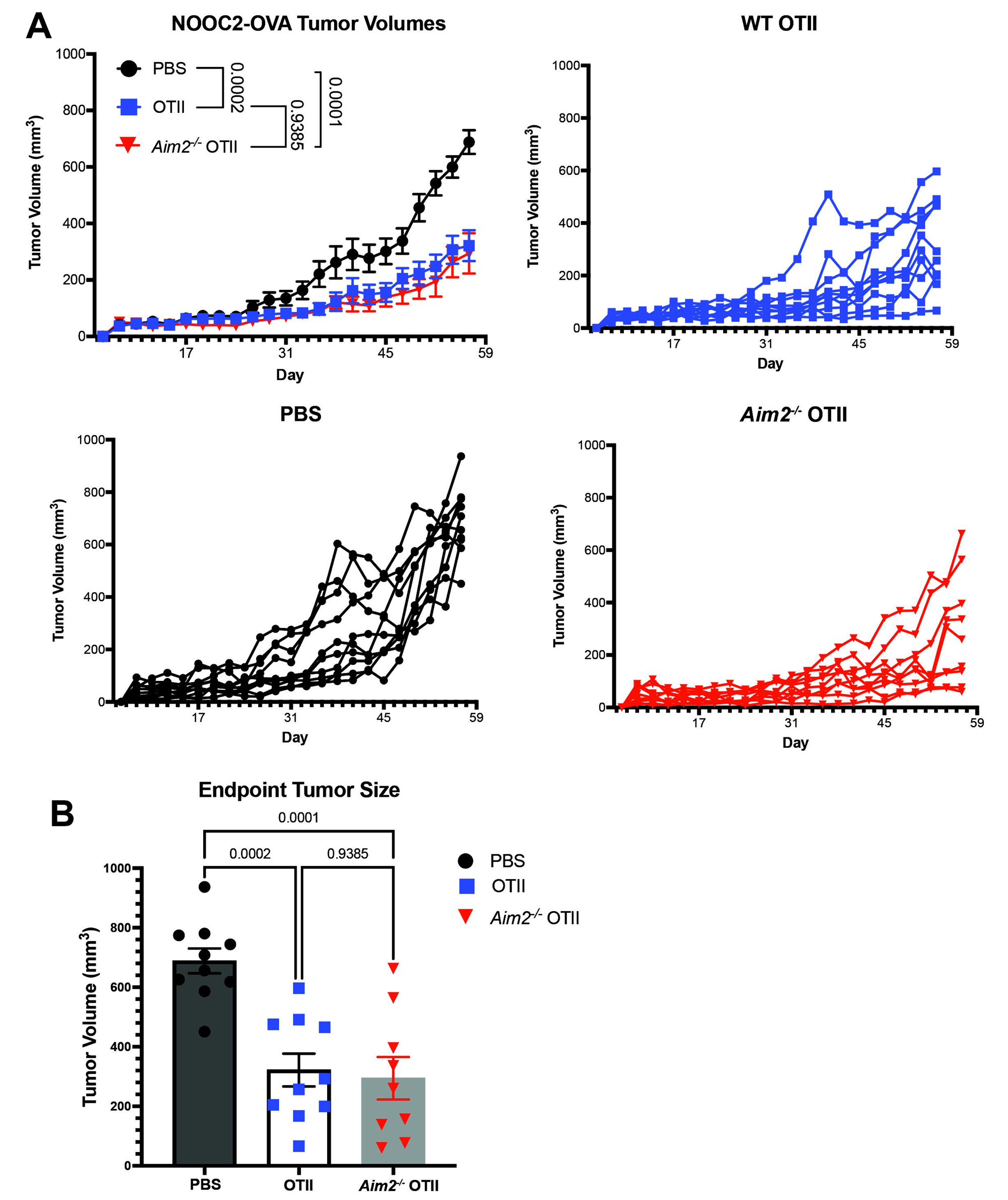
*Aim2^-/-^*OT-II CD4 T cells display no functional defects in promoting antigen-specific anti-tumor immunity. (**A**) NOOC2-OVA tumors were injected into C57BL/6J mice. At D10 post-injection, mice were adoptively transferred with either WT OT-II CD4^+^ T cells, *Aim2^-/-^* OT-II CD4^+^ T cells or PBS. (Top-left) Compiled tumor volumes of mice injected with PBS, OT-II or *Aim2^-/-^* OT-II CD4^+^ T cells. (Bottom-left) Tumor volumes for individual mice injected with PBS. (Top-right) Tumor volumes for individual mice injected with WT OT-II CD4^+^ T cells. (Bottom-right) Tumor volumes for individual mice injected with *Aim2^-/-^* OT-II CD4^+^ T cells. (**B**) Bar graph of NOOC2-OVA tumor volumes at the 56-day endpoint of panel A. Tumor volumes were measured by caliper. Data are presented as means +/-SEM and analyzed using a One-Way ANOVA with n=9-10 mice per group.

## Discussion

AIM2 is a PRR that exabits a growing list of diverse functions that impact immune responses. This includes its well described function in forming an inflammasome to release IL-1β and IL-18 during infection along with its ability to suppress AKT phosphorylation and modulate cellular metabolism. In the context of T cells, AIM2 can modulate the differentiation of CD4^+^ T cells into different subsets, including Tregs, Th17, and Tfh cells [10–13]. However, these studies depict conflicting roles for AIM2, possibly due to the use of different experimental disease models (e.g. EAE and IBD) and utilization of antibody-based in vitro skewing assays. In this study, we focused on the intrinsic role of AIM2 during antigen-specific activation of CD4^+^ T cells using well-established OVA-specific OT-II cell transgenic mice.

The data presented in this study contributes to our understanding of the function of AIM2 within and on CD4^+^ T cells during their development and antigen-specific activation, differentiation and effector function. Our findings indicate that AIM2 does not affect the overall number of CD4^+^ T cells that develop in the thymus, but it promotes positive selection of thymocytes and alters their maturation, migration, and survivability (or a combination of these factors) in the spleen. One explanation for these results is that *Aim2^-/-^* OT-II CD4^+^ T cells may emigrate to the spleen and mature faster than WT OT-II CD4^+^ T cells. This is supported by the fact that a greater proportion of *Aim2^-/-^* OT-II CD4^+^ T cells are Qa-2^high^, which is indicative a CD4^+^ T cell of being more mature after recent thymic emigration [19]. A potential molecular mechanism that could explain this may be rooted in the regulation of CD4^+^ T cell metabolism through AKT-mTOR signaling. Loss of AIM2 in activated T cells leads to overactive mTOR signaling [11], and suppression of the AKT-mTOR pathway is vital to maintaining quiescence and survival of naïve T cells in the spleen [28, 29].

Another possible explanation for the reduced numbers of splenic CD4^+^ T cells observed in AIM2 deficient mice is that AIM2 may promote thymic emigration. We observed that a larger percentage of thymocytes in *Aim2^-/-^* OT-II mice undergoing selection are CD69^+^, a marker that denotes T cells auditioning for selection. CD69 expression is upregulated very early after TCR stimulation and wanes once the stimulus ends [30]. While constitutive expression of CD69 on thymocytes in the thymus does not impair development, it does lead to the accumulation of mature SP thymocytes in the thymus and a reduction of T cells in the periphery [30]. This may also be caused by the excessive AKT signaling associated with AIM2 deficiency, thus resulting in constant low-level CD4^+^ T cell activation and maintenance of high levels of CD69 expression. Thus, it is possible that the thymocytes generated in *Aim2^-/-^* OT-II mice are unable to traffic out of the thymus resulting in the decreased number of CD4^+^ T cells observed in the spleens of these mice. Future experiments will determine if AIM2 controls CD4^+^ T cell emigration from the thymus and/or retention in the spleen during homeostatic conditions by regulating mTOR signaling and metabolic pathways in T cells.

IL-2 aids in T cell activation, expansion and Th1, Th2 and Treg cell differentiation. We revealed an intrinsic role for AIM2 in promoting IL-2 production and IL2ra expression in CD4^+^ T cells during antigen-specific activation in vitro. BLIMP-1 is a predominate transcription factor that suppresses IL-2 [31], however AIM2 did not impact BLIMP-1 expression in CD4^+^ T cells, suggesting this BLIMP-1-dependent negative feedback loop is likely not responsible for the reduced IL-2 during AIM2 deficiency. Alternatively, AIM2 may have a functional role in restricting the IL-2 negative feedback loop by suppressing AKT-mTOR signaling, which occurs downstream of TCR and CD28 signaling [32, 33]. IL-2 signals through the Akt-mTOR pathway [22, 24], which also initiates the IL-2 negative feedback loop. This suggests AIM2 deficient naïve T cells may be more sensitive to IL-2 signaling and therefore require less IL-2 for their activation and division. If correct, the IL-2 negative feedback loop may be triggered early when AIM2 is absent, which could result in early termination of IL-2 production. This notion is supported by the findings that *Aim2^-/-^* OT-II CD4^+^ T cells display decreased IL-2 production in vitro, but these cells proliferated and differentiated normally upon antigen challenge in vivo.

Although loss of AIM2 in OT-II CD4^+^ T cells resulted in a dramatic reduction in IL-2 production in response to antigen presentation in vitro, we failed to detect any functional defects in T cell function during primary CFA-OVA immunization or OVA-NOOC2 tumor challenge in vivo. This may be due to the presence of other tissue cytokines and DAMPs that promote T cell survival and differentiation, such as IL-7 and ATP [6, 34]. This is particularly important during disease states with distinct DAMP, cytokine and microbial environments (e.g., autoimmune encephalitis vs. colitis) where AIM2 has been implicated in driving T cell differentiation [10–12] and could help to explain the discrepancies between these and our findings. However, IL-2 signaling is also crucial to mounting an appropriate immune response to secondary challenge through the development of CD4^+^ T cell memory and expansion of CD8^+^ memory T cells [35, 36]. Furthermore, IL-2 is also required for the generation of viral-specific CD4^+^ Th1 tissue resident memory cells in the lung [37]. Therefore, while AIM2 does not appear to impact primary CD4^+^ T cell responses, it may play a more predominate and detectable role during antigen-specific rechallenge through IL-2-driven memory T cell generation. DAMPs are important contributing factors that can drive memory T cell formation [6]. As AIM2 can act as a DAMP sensor for self dsDNA, its DAMP-sensing function may also contribute to regulating memory formation in combination and/or distinct from its ability to regulate IL-2.

In conclusion, we showed that AIM2 contributes to CD4^+^ T cell homeostasis in the secondary lymphoid organs as evidenced by reduced numbers CD4^+^ cells and a reduction of recent thymic emigrants in the spleen of *Aim2^-/-^* OT-II mice. We also show AIM2 intrinsically promotes IL-2 production from OT-II CD4^+^ T cells in response to antigen-specific activation in vitro. Surprisingly, this loss of IL-2 in vitro did not translate into measurable differences in OT-II CD4^+^ T cell proliferation or differentiation in response to OVA immunization in vivo. Moreover, AIM2 was dispensable for OT-II CD4^+^ T cell effector function during challenge with syngeneic OVA-expressing tumors. Because our experiments only utilized primary challenge models, future studies will also focus on the role of AIM2 during memory T cell development during antigen-specific infection models and if this is linked to AIM2’s PRR function in sensing DAMPs.

## Acknowledgments

The authors would like to thank Yu Leo Lei, (MD Anderson) for generating and providing the NOOC2-OVA cell line. The authors would also like to thank the University of Arizona Cancer Center (UACC) Experimental Mouse Shared Resource (EMSR) and the Flow Cytometry and Immune Monitoring Shared Resource for equipment and technical assistance. Research support proved by UACC shared resources was supported by the National Cancer Institute of the National Institutes of Health under award number **P30 CA023074.** The authors also thank Sydney A.N. Verdugo and Anika M. Arias for their helpful editorial comments.

## Footnotes

This work was supported by an Arizona Biomedical Research Center New Investigator Award CTR056057 (J.E.W), National Institutes of Health, National Institute of Diabetes and Digestive and Kidney Diseases Grant R01DK141491 (to J.E.W.), the National Institutes of Health, National Institute of Dental Craniofacial Research Grants F31DE032263 (to D.M.R), R01DE026728 (to Y.L.L) and U01DE033330 (to Y.L.L), the National Institutes of Health, National Institute on Aging Grant T32AG058503 (to D.M.R) and the National Institutes of Health, National Institute of Allergy and Infectious Diseases Grants R01AI101053 (to M.S.K) and R01AI179713 (to M.S.K).

